# Investigating Gene Expression Noise Reduction by MicroRNAs and MiRISC Reinforcement by Self-Feedback Regulation of mRNA Degradation

**DOI:** 10.1101/2025.02.11.637731

**Authors:** Shuangmei Tian, Ziyu Zhao, Meharie G. Kassie, Fangyuan Zhang, Beibei Ren, Degeng Wang

**Affiliations:** Department of Environmental Toxicology, and The Institute of Environmental and Human Health (TIEHH), Texas Tech University, Lubbock, TX 79416, USA; Department of Mathematics and Statistics, Texas Tech University, Lubbock, TX 79409-1042, USA; Department of Mechanical Engineering, Texas Tech University, Lubbock, TX 79409-1021, USA

## Abstract

The microRNA (miRNA) induced silencing complex (miRISC) is the targeting apparatus and arguably the rate-limiting step of the miRNA-mediated regulatory subsystem – a major noise reducing, though metabolically costly, mechanism. Recently, we reported that miRISC channels miRNA-mediated regulatory activity back onto its own mRNAs to form negative self-feedback loops, a noise-reduction technique in engineering and synthetic/systems biology. In this paper, our mathematical modeling predicts that mRNA expression noise exhibits a negative correlation with the degradation rate (K_deg_) and is attenuated by self-feedback control of degradation. We also calculated K_deg_ and expression noise of mRNAs detected in a total-RNA single-cell RNA-seq (scRNA-seq) dataset. As predicted, miRNA-targeted mRNAs exhibited higher K_deg_ values accompanied by reduced inter-cell expression noise, confirming the operational trade-off between noise suppression and the increased metabolic/energetic costs associated with producing these mRNAs subjected to accelerated degradation and translational inhibition. Moreover, consistent with the K_deg_ self-feedback control model, miRISC mRNAs (AGO1/2/3 and TNRC6A/B/C) exhibited further reduced expression noise. In summary, mathematical-modeling and total-RNA scRNA-seq data-analyses provide evidence that negative self-feedback regulation of mRNA degradation reinforces miRISC, the core machinery of the miRNA-mediated noise-reduction subsystem. To our knowledge, this is the first study to concurrently analyze mRNA degradation dynamics and expression noise, and to demonstrate noise reduction by self-feedback regulation of mRNA degradation.

## Introduction

MicroRNA (miRNA) mediates a gene expression regulatory subsystem that functions throughout the plant and animal kingdoms, though the animal and plant sub-systems are very different from each other^1-4^. In animals, miRNA genes can be either independent transcription units (TU) or part of the introns of other genes^1,5,6^. A miRNA gene may encode a single miRNA hairpin precursor, or a cluster of multiple precursors. In canonical miRNA biogenesis pathway, the primary transcript is cleaved into the precursor hairpins that, upon nucleus export into the cytoplasm, are further processed by Dicer and loaded into the Argonaute (AGO) proteins. In non-canonical pathways, one or more steps are skipped^7-11^. AGO proteins then remove the complementary strand of the double-stranded precursor to form the mature miRNAs. The miRNA-AGO complex binds to cognate miRNA binding sites, usually in the 3’-UTR of the target mRNAs^12^, leading to moderate translation inhibition, degradation enhancement or both.

The miRNA regulatory subsystem has three segments: 1) biogenesis of mature 22-nucleotide miRNAs described above; 2) formation of the miRISC targeting apparatus, arguably the rate-limiting and thus the most critical segment of this mRNA regulation subsystem; and 3) effectors for mRNA decay and translation inhibition. MiRISC consists of the miRNA-loaded AGO protein and AGO-recruited TNRC6 protein upon miRNA binding to cognate sites on target mRNAs. The complex channels moderate inhibitory actions onto target mRNAs via TNRC6-recruitment of the effectors, with a single miRNA binding site conferring only ∼20% inhibition.

Why do the cells maintain this regulatory subsystem that seemingly violates the cellular gene expression economics? It is wasteful to expend critical metabolic and energetic resources to produce the miRNA-targeted mRNAs but then render them translationally inhibited and under enhanced degradation pressure^13^. What are the operational advantages the cells gain as a trade-off for the wasted metabolic and energetic resources?

One such advantage is the well-documented miRNA-mediated reduction of cell-to-cell gene expression fluctuation, often termed noise, such as those incurred by the stochastic transcriptional bursting^14-16^, thereby enhancing robustness of cellular processes^17^. That is, expression levels of targeted mRNAs and their proteins have reduced cell-to-cell fluctuations, enhancing homeostasis and cellular homogeneity. The importance of cellular homeostasis and homogeneity is exemplified by their disruption in cancers to enable cellular adaptability to spatiotemporally unpredictable environments, leading to uncontrollable clonal growth and metastasis^18^. However, the noise-reduction capacity of the sub-system is limited when acting alone, *i*.*e*., open-loop regulation^19^. Consequently, miRNAs often partner with other regulators, especially transcription factors (TF), to implement common system control strategies such as feedback, known as closed-loop control^20^.

We previously reported that miRISC is itself controlled directly by miRNAs to form a negative self-feedback loop^21,22^ – a proven noise-reduction technique in engineering and synthetic/systems biology^23-27^. That is, the core miRISC apparatus is controlled by a negative closed-loop, while other miRNA-targeted mRNAs by either open loops or closed loops formed with other regulators. We looked for opportunities to investigate whether and how this self-feedback loop complements basic miRNA regulatory activity to reinforce the stability of miRISC mRNA expression.

Single-cell RNA-seq (scRNA-seq) is a powerful extension of the bulk RNA-seq technology that gains the capacity to analyze cell-to-cell gene expression noise, and cellular sub-populations^28,29^. ScRNA-seq methods can be roughly divided into two categories. First, the traditional scRNA-seq methods, exemplified by the 10x Chromium methods^30^, are typically microfluidic droplet-based and sequence the ends of RNA molecules. They are capable of economical high, and still increasing, throughput analyses, currently tens of thousands of cells in one experiment. But they have low gene detection sensitivity^29^. Low expression level mRNAs, which unfortunately include many regulatory genes such as TFs, are likely to evade their detection in most of the analyzed cells. Second, thus, a new category of methods recently emerged to complement them^31-34^. The new methods enhance gene detection sensitivity by covering full length RNAs. They are often plate based and have much lower throughput, currently analyzing hundreds of cells in an experiment. Successful applications of the methods to analyze miRNA-mediated regulatory activity, such as gene expression noise reduction, have been reported^35-37^. Additionally, many methods have a total RNA version, that is, simultaneously analyzing the transcriptome at nascent intron-containing and mature fully spliced RNA steps. To the best of our knowledge, total RNA scRNA-seq datasets have not been explored to advance our understanding of miRNA-mediated regulatory activities.

Thus, in this study, we explored a dataset of snapTotal-seq, the latest total-RNA scRNA-seq method capable of detecting over 10,000 genes at both nascent and mature RNA steps in a cell^34^. MiRNA-targeted mRNAs exhibited low expression noise in conjunction with enhanced degradation. We also observed a further reduction of expression noises of AGO1/2/3 and TNRC6A/B/C mRNAs, supporting the role of the self-feedback loop in reinforcing miRISC – the core of the miRNA-mediated noise reduction subsystem. The same was observed for the RNA binding protein QKI that was recently reported to be an auxiliary partner for miRNA-AGO regulatory activity^38^; QKI is also regulated by miRNAs^21,22^ and, thus, an auxiliary component of the miRISC negative self-feedback loop. Additionally, AGO1/2/3 mRNA expression noise reduction occurred at the mature mRNA step, whereas TNRC6A/B/C noise reduction occurred at the nascent RNA step and was further reinforced at the mature mRNA step.

## Materials and Methods

### SnapTotal-seq dataset and analysis

The snapTotal-seq dataset is publicly available at the NCBI GEO database under the accession number GSE202126. We downloaded the dataset and processed it following the procedure described in the original publication^34^. Briefly, the next generation sequencing (NGS) reads were mapped to human genome assembly (GRCh37) with the STAR aligner (version 2.5.3a). Then, uniquely mapped reads were mapped to the GENCODE gene annotation (version 19) with the htseq-count software. Nascent- and mature-RNA originated reads were distinguished by the presence of intronic sequences. To normalize nascent and mature RNA expression levels (read counts) separately with respective total read counts of each cell, we calculated the counts for each gene per 100,000 mapped reads. The normalized read counts were used in downstream analysis. For details, please see the original publication^34^.

### Quantification of nascent and mature RNA inter-cell expression level noise

We used the coefficient of variation (CV) parameter to quantify inter-cell RNA expression noise, as CV has been routinely used for such purposes^19,35,36^. We examined the cross-cell distributions of nascent and mature RNA expression levels to determine how to calculate this parameter.

For mature mRNAs, we observed that the normalized read counts of individual genes across the cells already resembled normal distributions. So, we pre-process the read counts with the cell-cycle scoring and regression function in the SEURAT single cell genomics R tool kit^39^, mitigating the effect of cell cycle heterogeneity. We then calculated the standard deviation (*σ*) (SD) and the mean (*μ*) for each gene, and then CV_mat_ (mat: mature) as the SD-to-mean ratio:

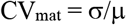

At the nascent RNA step, due to stochastic transcriptional bursting, RNA copy-numbers across a cell population usually follow the Poisson distribution or Poisson distribution with zero spikes, also known as zero-inflated Poisson distribution^14-16^. Consistently, we observed that the histogram of normalized read counts for each gene across the cells resembles a log-normal (LN) distribution. We thus log_2_-transformed the counts, pre-processed the data to mitigate the effect of cell cycle heterogeneity, and calculated the SD (*σ*) and mean (*μ*) of the transformed data for each gene. The normalized read counts across the cells can then be notated as LN(*μ, σ*^2^). The CV_nas_ (nas: nascent) of the original LN distribution can then be calculated in the standard procedure as below:

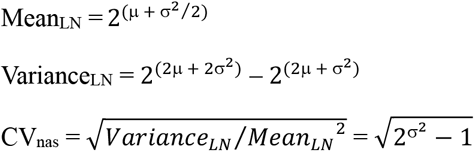

### Mathematical modeling of mRNA expression and inter-cell noise without feedback regulation

In the biomedical literature, mRNA expression is usually modeled as a first-order dynamic process to describe the relationship among mRNA expression level (R), production rate (P), degradation rate (K_deg_), steady-state expression level (R*) and inter-cell expression level stability (S)^23,40^; in engineering, such a process is referred to as “first-order system”. When no feedback is involved, the model (*f*_unreg_(R), unreg: not self-feedback regulated) is shown below:

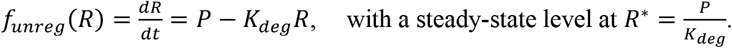

To experimentally determine the mRNA production rate P, multiple approaches have been developed. Since RNA splicing co-occurs with and is faster than transcription^41-43^, P can be approximated as the transcription rate, which is measured by the GRO-seq technology. Alternatively, P may be expressed as the product of nascent RNA expression level (R_nas_) and the nascent RNA splicing rate (K_sp_), simplified as K_sp_R_nas_^40,44^. Recently, this formulation has served as the basis of RNA velocity analysis facilitated by total-RNA scRNA-seq methods^33,34,40^. Thus, as in many other studies^33,34,40,44^, we used K_sp_R_nas_ in this study as it was measured by snapTotal-seq. Thus, for mRNAs at steady-state level,

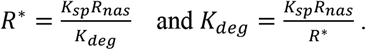

The expression level R transits from a non-steady-state level R^0^ to R^*^ as shown below:

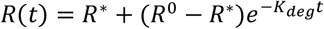

K_deg_ quantifies the speed of the R^0^-to-R^*^ transition and, thus, reflects the mRNA expression-level stability without feedback regulation (S_unreg_); in engineering, 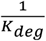 is defined as the system time constant. Alternatively, S_unreg_ can be calculated as the noise decay rate via Taylor expansion, yielding the same result^23^:

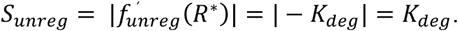

Thus, a higher K_deg_ predicts higher expression level stability and, thus, lower noise (CV).

### Modeling of mRNA expression and inter-cell noise with self-feedback regulation

The self-feedback loop adds a regulatory element to the degradation rate (K_deg_), highlighted by the red bold text in the model (*f*_reg_(R), reg: self-feedback regulated) below:

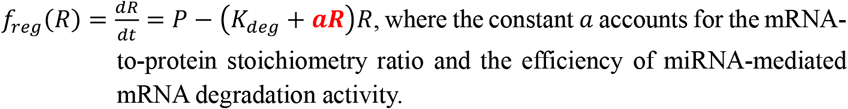

Taylor expansion also allows calculation of mRNA expression-level stability with self-feedback regulation (S_reg_), and the regulated-to-unregulated stability ratio (S-ratio) as below^23^:

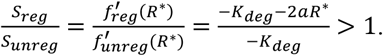

Thus, the self-feedback loop is expected to enhance inter-cell mRNA expression level stability (S_reg_ > S_unreg_) and consequently reduce inter-cell noise (CV_reg_ < CV_unreg_).

### Statistical analysis

The R open-source software (version 4.0.2) was used for data analysis and plotting. Standard R functions were used for sample normalization, parameter calculation, linear regression and LOESS regression.

For a weighted linear regression, we first performed an un-weighted linear regression. Then the reciprocals of the square of the residuals were used as the weights to perform the weighted linear regressions. The weighted residuals of the weighted regression were used for subsequent analysis.

To compare the linear regression models, we used the ANOVA function, as one model is nested within the other. The function outputs an F-ratio and a p-value, thus statistically quantifying the improvement from adding the second predictive variable.

## Results

### Different gene expression patterns at the transcription/nascent-RNA and mature RNA steps due to post-transcriptional regulation

In this study, we used the terms “transcription” and “nascent RNA” as inter-exchangeable terms to denote the gene expression step. Nascent RNA expression level R_nas_ is determined by transcription rate (TR) and nascent RNA splicing rate (K_sp_) and, at steady state, equal to the TR/K_sp_ ratio. R_nas_ and TR are thus frequently used in the literature as a measure of each other. As described in the section of Materials and Methods, we downloaded the snapTotal-seq dataset from the NCBI GEO database, mapped the reads onto the human genome and used the presence of intron sequences to distinguish nascent RNAs from their mature RNA counterparts. We then normalized the nascent and mature RNA read counts across the cells by calculating counts per 100,000 mapped reads for each gene.

It has been reported by us that gene expression is less selective at the transcription step than the mature mRNA step^44^. We tested this pattern with genes detected at both nascent RNA and mature mRNA steps in every cell in the snapTotal-seq dataset (Fig. 1). A comparative boxplot shows that the normalized read counts exhibited a narrower value range and reduced dispersion at the nascent RNA step compared to the mature spliced RNA step (Fig. 1A). That is, at the nascent RNA step, the gene expression levels were more uniform. In contrast, at the mature mRNA step, a portion of the genes were expressed at much higher levels, presumably to meet near-term cellular protein production needs, whereas another portion of the genes were expressed at much lower levels.

**Figure 1.**
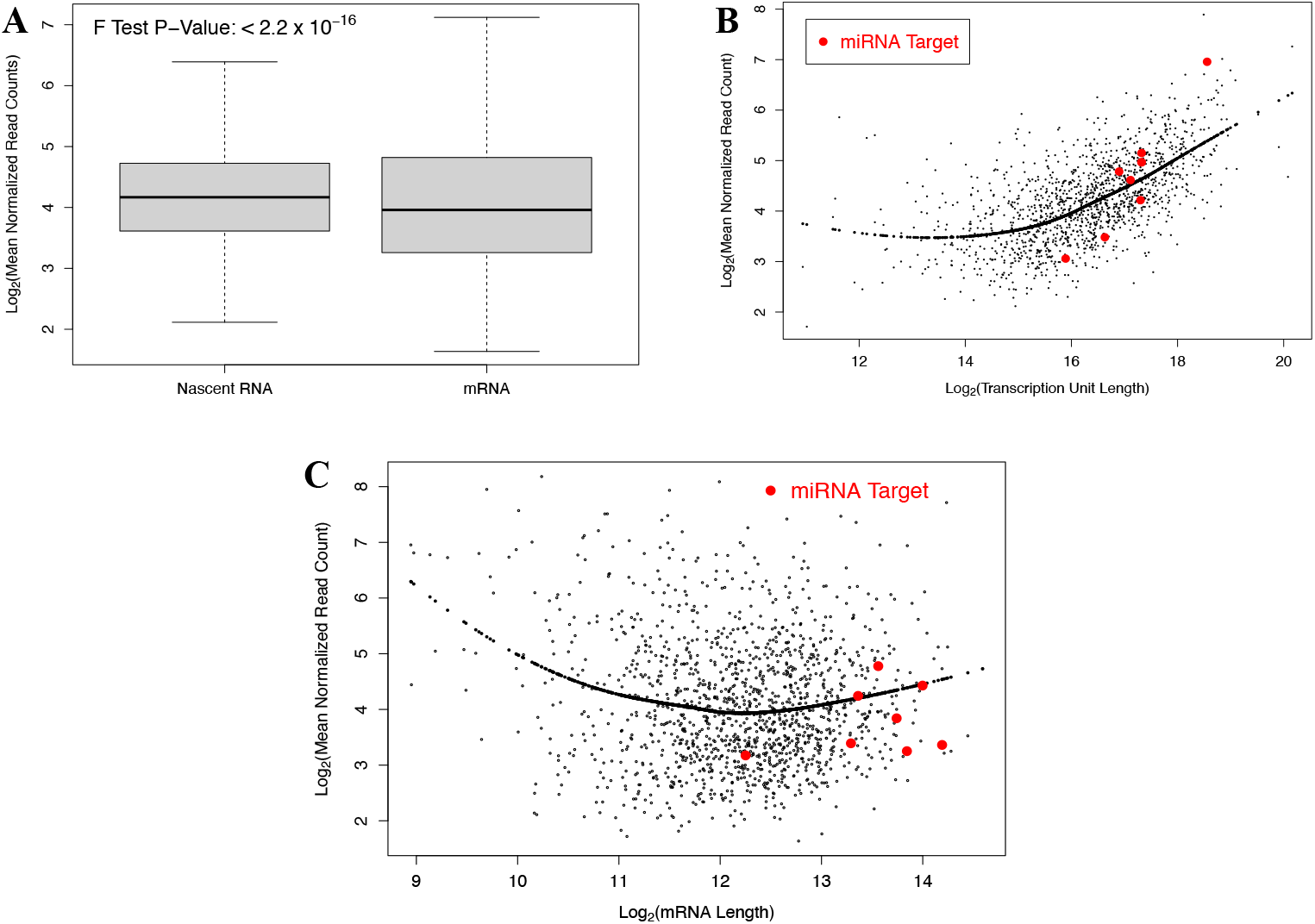
Change of expression patterns from nascent to mature RNA steps. Comparative analyses of nascent and mature RNA expressions are shown. Genes detected at both steps in all cells were used. **A:** A comparative boxplot of nascent/pre-splicing and mature RNA expression levels shows enhanced selectivity or dispersion level at the mature RNA step. **B:** A scatter plot of nascent RNA expression level versus transcription unit (TU) length shows positive correlation between the two. A LOESS regression curve is added to illustrate the relationship. The nascent RNAs whose mature RNA counterparts have >= 60 miRNA binding sites are highlighted. **C:** A scatter plot of mature mRNA expression level versus length is shown. A LOESS regression curve is added to illustrate the loss of the correlation observed in **B.** The mRNAs with >= 60 miRNA binding sites are highlighted.

However, the normalized read count data missed a key factor – RNA length. Since the snapTotal-seq sequencing chemistry covers full-length RNA, the length becomes one of the determining factors for the read counts. The NGS read counts (K) are known to resemble the Poisson distribution:

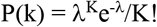

For individual RNAs, the means (*λ*) of the distributions should be directly proportional to their lengths. Consistently, snapTotal-seq gene detection rates were higher for longer RNAs at both nascent and mature RNA steps^34^.

RNA length gave us another opportunity to test the lower gene expression selectivity at the transcription step. The lower selectivity should make RNA length a more prominent factor for the read counts at this step than the mature RNA step. Not surprisingly, we observed a significant positive correlation between the nascent RNA read count and the transcription unit (TU) length – the length of the longest possible un-spliced RNA of a gene (Fig. 1B). However, the correlation essentially disappeared at the mature RNA step (Fig. 1C). Conceivably, additional regulatory actions on RNA splicing, nucleus export and RNA degradation led to higher gene expression selectivity at mature RNA step, obscuring the impact of RNA length.

### Adjusting the read counts with RNA length

Nevertheless, the full-length RNA coverage by the snapTotal-seq sequencing chemistry imposed a need to adjust the read count data by RNA length, so that the read counts of RNAs of different lengths were comparable with one another. The read count of a RNA is determined by both its expression level, *i*.*e*., copy number, and its length. With everything else remaining the same, the expected read count of a RNA, *i*.*e*., the mean (*λ*) of the Poisson distribution, should be, as discussed above, directly proportional to the RNA length.

Given the obvious correlation between nascent RNA read count and TU length (Fig. 1B), we adjusted the nascent read count with a LOESS regression (log_2_(read-counts) versus log_2_(TU-length)). Consequently, the residuals of the regression were used as the adjusted read counts. Regarding the mature RNA step, due to the disappearance of the correlation, we used the read counts normalized by RNA length – the RPKM value popularly used in RNA-seq data analysis.

### Mathematical model of gene expression noise

As discussed in the Materials and Methods section, we followed the conventional practice of modeling the mature mRNA expression without feedback control as a first-order dynamic process and derived relationship among the expression parameters. The model predicts that higher degradation activity, *i*.*e*., the K_deg_ value, should lead to higher steady-state expression level (R*) stability (S_unreg_). CV_unreg_ that quantifies inter-cell fluctuation of R* is the reciprocal of S_unreg_ as shown below:

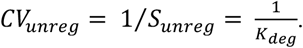

Thus, higher K_deg_ predicts higher expression level stability and thus lower CV values^23,40^. We explored the snapTotal-seq dataset to estimate CV and K_deg_ and to test predicted relationship between the two.

### Analyzing inter-cell gene expression noise

The inter-cell fluctuation of the gene expression levels has two contributing factors. The first is the gene expression noise due to stochasticity of the gene expression process – the target of this study. The second is the noise intrinsic to experimental procedures, which the NGS library preparation and sequencing processes are known to have. It is also known that noise from the second source is higher for low-expression-level RNAs than high-expression-level RNAs. As shown in figure 2A, the difference between the mature mRNA read counts of the same genes in two cells decreases as the expression level increases. And the same trend was observed for the nascent RNA read counts (Fig. 2B). Therefore, we analyzed the inter-cell RNA expression noises in the context of mean expression levels. As shown in figures 2C and 2D, the CV value decreases linearly as the mean expression level increases at both the mature mRNA and the nascent RNA steps, with the mature mRNA step having a steeper slope than the nascent step.

**Figure 2.**
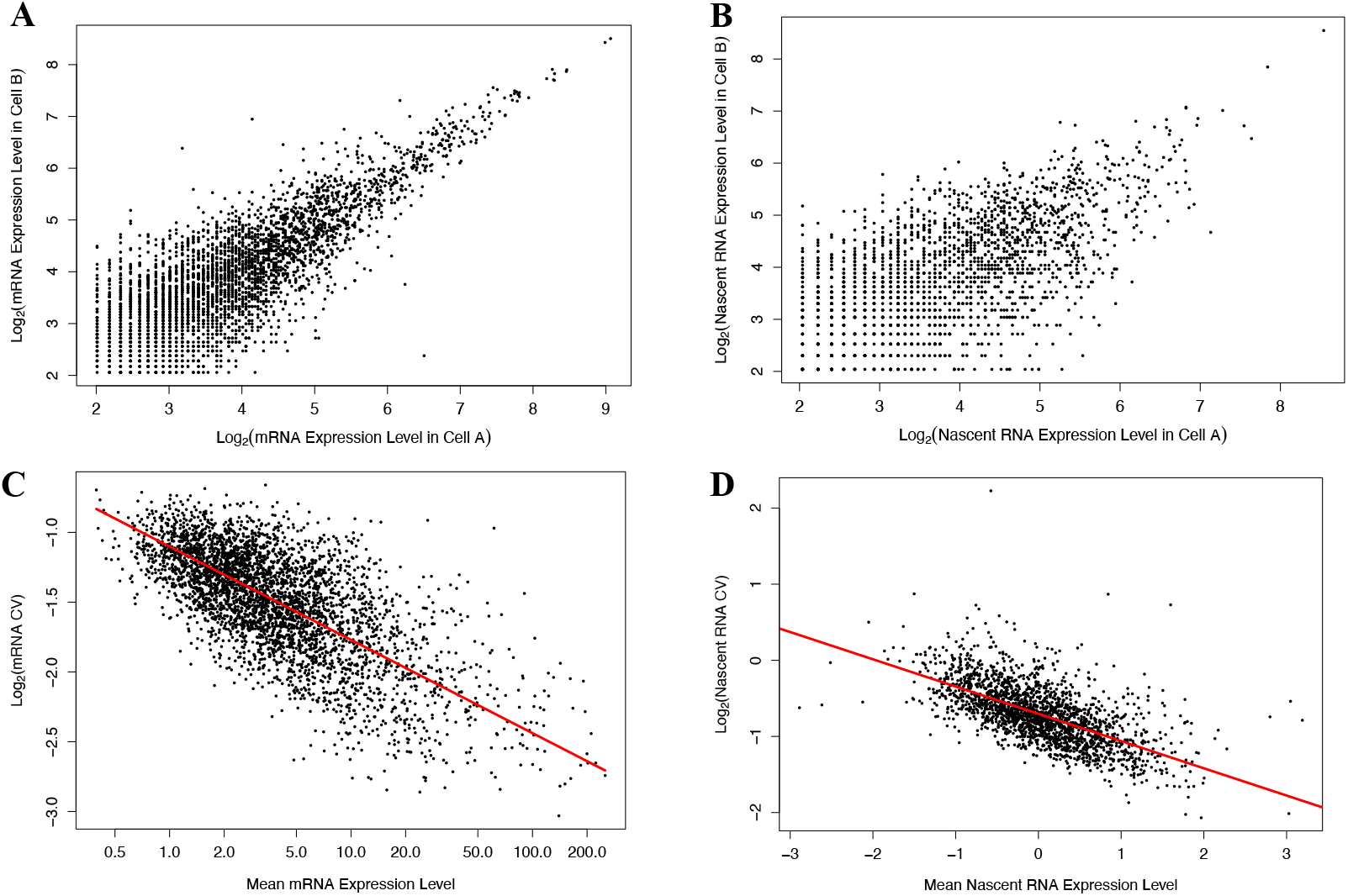
Analyzing noises observed in the snapTotal-Seq dataset. **A and B:** comparative scatter plots of RNA expression levels at the mature mRNA (**A**) and the nascent RNA (**B**) steps in the same pair of cells, illustrating a pattern of decreasing inter-cell fluctuation as expression level increases at both steps. **C and D:** scatter plot of RNA CV versus mean expression level at the mature mRNA (**C**) and the nascent RNA (**D**) steps. The pattern of decreasing inter-cell fluctuation, which is quantified by CV, as mean expression level increases is illustrated at both steps. Weighted linear regression line is shown to illustrate the pattern. CV: coefficient of variation.

A comparative analysis of nascent and mature RNA expression noise was then performed (Fig. 3). Weighted linear regressions were conducted (CV versus expression levels), with the regression lines shown in figures 2C and 2D. We used the residuals of the regressions as adjusted CV and then plotted the scaled adjusted CV in a scatter plot (Fig. 3). As expected, a positive correlation was observed (Fig. 3). However, the correlation was moderate, with a correlation coefficient of 0.315, and the slope of the linear regression is only 0.33. Thus, post-transcriptional regulatory mechanisms must have reduced the impact of propagation of nascent RNA expression noise on the mature RNA expression noise. Otherwise, the correlation should be much stronger, and the linear regression slope should be much closer to 1.

**Figure 3.**
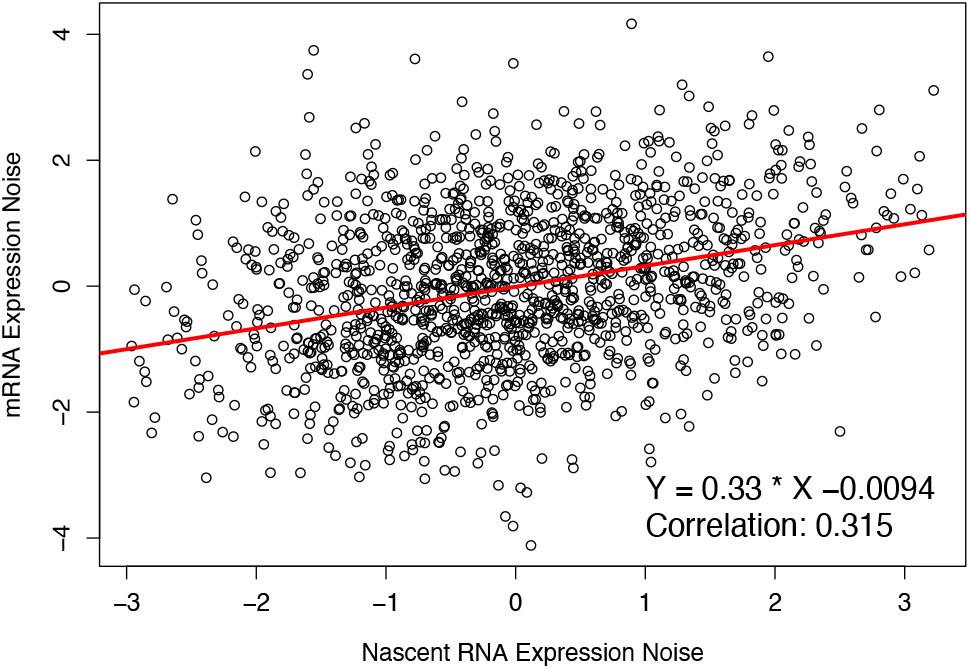
Scatter plot of inter-cell expression level fluctuations of mature mRNAs versus those of nascent RNAs. Mean expression level adjusted CV values, *i*.*e*., the residuals of the linear regressions in figures 2C and 2D, were used for the plot. A linear regression line is shown. The equation of the linear regression and the correlation coefficient of the two variables are also shown.

### Relationship between mRNA degradation activity and inter-cell expression noise

Having both nascent and mature RNA expression levels simultaneously allowed us to test our mathematical prediction. The nascent RNA expression levels were used in the literature in lieu of transcription and/or mRNA production rates in mRNA expression modeling. Therefore, we estimated the mature RNA K_deg_ as the mature to nascent RNA expression level ratio (see Materials and Methods). However, there are intrinsic complexities that contribute to mature RNA expression noise observed in scRNA-seq datasets. The propagation of nascent RNA expression noise, which variated from gene to gene, was discussed above (Fig. 3).

We tested whether higher degradation activity led to reduced inter-cell mature RNA expression noise and whether the trend was strong enough to overcome these interfering factors. Our approach was a linear regression of the data shown in figure 2C with K_deg_ added as the second predictive variable. The result is schematically shown in figure 4A. Despite these complexities, the analysis revealed a clear trend of decreasing CV values along increasing K_deg_ values. ANOVA comparison of the regression models determined that adding K_deg_ as the second predictive variable led to significant improvement (F_1_ = 16.8, p-value = 4.37E-5).

**Figure 4.**
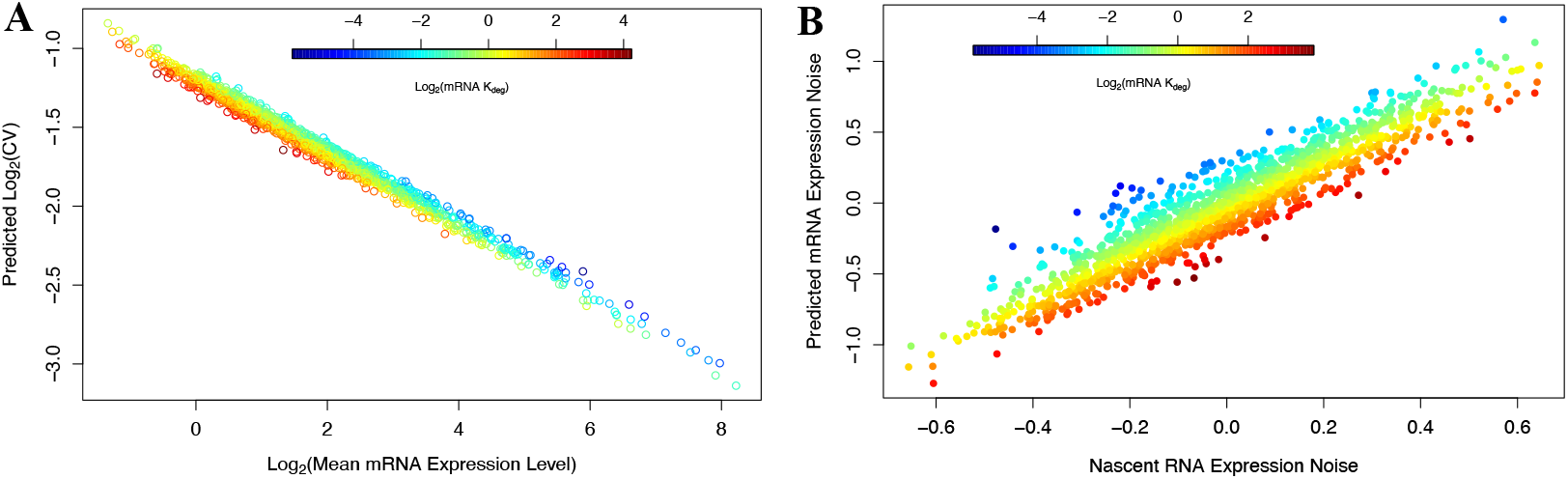
High degradation activity tends to reduce mature mRNA expression noise. **A:** Linear regression analysis of data shown in figure 2C, with mRNA K_deg_ added as the second predictive variable (CV versus (mean expression level) + K_deg_). Predicted CV values are plotted versus the mean expression levels, with the data points color coded by K_deg_. **B:** Linear regression analysis of data shown in figure 3, with mRNA K_deg_ added as the second predictive variable (CV_mat_ versus CV_nas_ + log2(K_deg_)). Predicted CV_mat_ values are plotted versus the CV_nas_ values, with the data points color coded by K_deg_. Mat: mature. Nas: nascent.

Subsequently, we tested this trend via a comparative analysis of expression-level adjusted nascent and mature RNA CV values that, as shown in figure 3, had a moderately positive correlation with each other. Our approach was a linear regression with K_deg_ added as the second predictive variable. The result is schematically shown in figure 4B. The regression predicted a clear trend that mature RNAs with high K_deg_ values tended to have lower CV values relative to nascent RNAs (Fig. 4B). ANOVA comparison of the regression models determined that adding K_deg_ as the second predictive variable led to significant improvement (F_1_ = 12.4, p-value = 0.00045).

Taken together, these results supported the notion that mature RNA degradation activity exerted prominent impact on inter-cell mature RNA expression noise, validating our mathematical model.

### Reduced expression levels and inter-cell noises of miRNA-targeted mRNAs

Encouraged by these observations, we next explored whether miRNA-mediated target mRNA degradation and inter-cell expression noise reduction can be simultaneously assessed. Performing such analysis with traditional high-throughput scRNA-seq has been technically challenging due to the low sensitivity, *i*.*e*., low gene detection rate, of these scRNA-seq methods. MiRNA-targeted mRNAs are usually not expressed at high levels, thus evading detection by these methods. SnapTotal-seq substantially alleviates this low sensitivity issue. And it exhibits higher gene detection rate for long RNAs^34^. It is thus well-suited for studying regulatory activity mediated by miRNAs, whose target mRNAs also tend to be longer.

We calculated the count of unique evolutionarily conserved miRNA binding sites for each mRNA. Indeed, most top miRNA-targeted mRNAs, *i*.*e*., those with highest conserved binding site counts, were detected by this scRNA-seq method. We identified 23 mRNAs with 60 or more conserved binding sites. 16 of them were detected in every cell at the mRNA step, and 10 of them at the nascent RNA step. 8 of them were detected in every cell at both steps. As shown in figure 1C, the 8 miRNA-target mRNAs exhibited lower-than-expected expression levels. The pattern was specific for the mature mRNA step. At nascent RNA step, their counterpart RNAs exhibited normal expression levels (Fig. 1B).

To simultaneously analyze miRNA-mediated target mRNA degradation and expression noise reduction, we re-plotted the mature mRNA CV values versus length-adjusted mean expression level shown in figure 2C, but with the data points color-coded by the miRNA binding site counts (Fig. 5A). The plot revealed that mRNA expression levels decreased in a binding site count dependent manner, and so did the inter-cell expression noise. Both trends were evident without the need for regression analysis (Fig. 5A).

**Figure 5.**
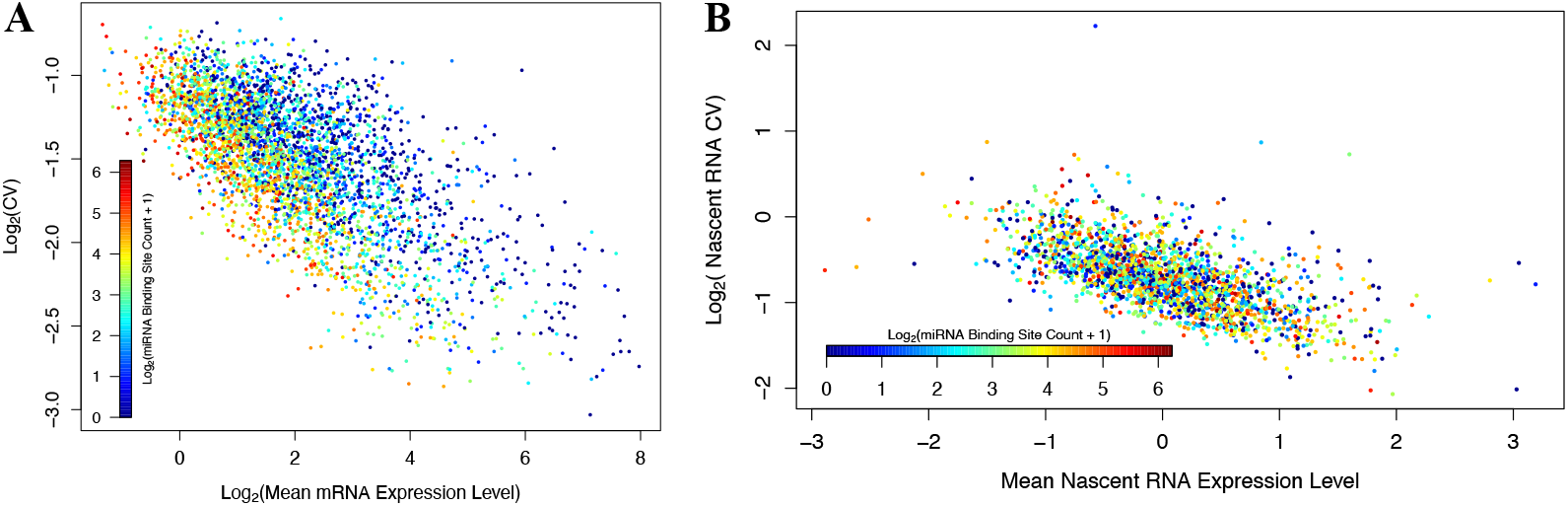
MiRNA targeted mRNAs have lower expression levels and expression noises, but their nascent RNA counterparts do not. **A:** Re-plot of figure 2C, with the data points color coded by miRNA binding site counts. **B:** Re-plot of figure 2D, with the data points color coded by miRNA binding site counts.

We directly analyzed miRNA-mediated target mRNA degradation via a comparative analysis of nascent and mature RNA expression levels. Our approach was a linear regression with log_2_(miRNA binding site count) added as the second predictive variable. The result is schematically shown in figure 6A. The regression predicted a clear trend that mature RNAs with high binding site counts tended to have lower expression levels compared to nascent RNAs (Fig. 6A). ANOVA comparison of the regression models demonstrated a significant

**Figure 6.**
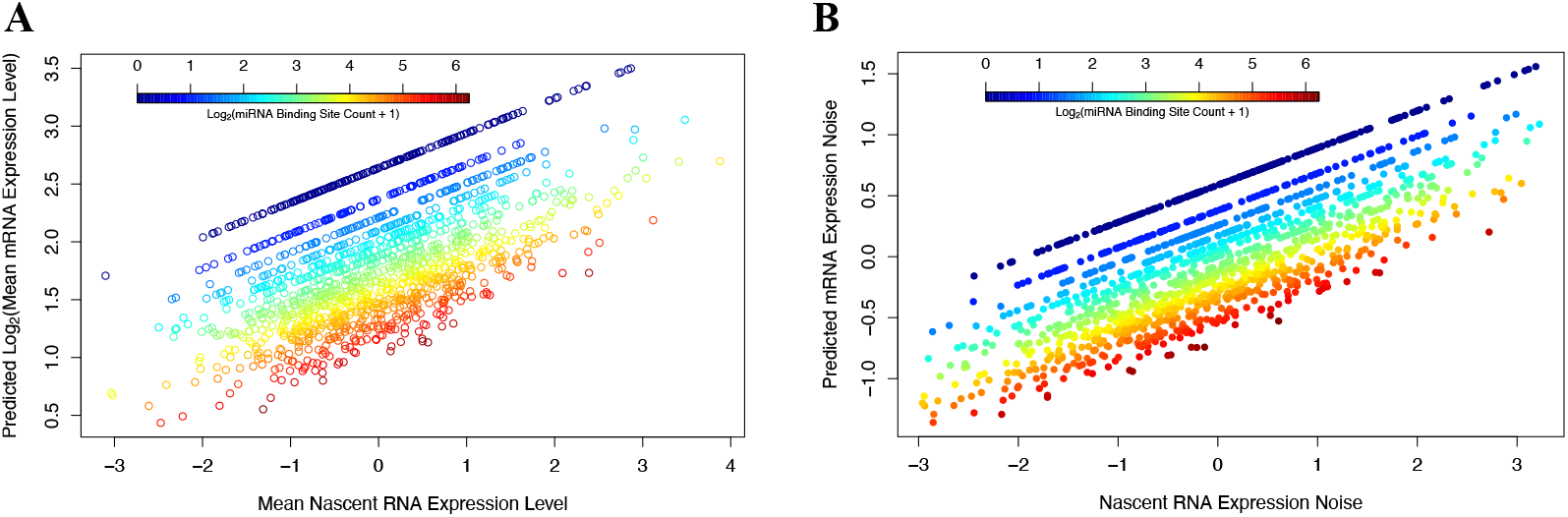
Impact of miRNA-mediated regulation on target mRNA expression levels **(A)** and noise **(B)** revealed by linear regression analyses. **A:** Mature mRNA expression level versus nascent RNA expression level linear regression, with miRNA binding site count added as the second predictive variable. Predicted mature mRNA expression level was plotted versus the nascent RNA expression level, with the data points color coded by miRNA binding site counts. **B:** Linear regression analysis of data shown in figure 3, with miRNA binding site count added as the second predictive variable (CV_mat_ versus CV_nas_ + log_2_(Site Count)). Predicted CV_mat_ values are plotted versus the CV_nas_ values, with the data points color-coded by miRNA binding site count. improvement in the predictive power of the model when including the second predictive variable (F_1_ = 143.34, p-value < 2.2E-16).

We also analyzed miRNA-mediated target mRNA expression noise reduction via a comparative analysis of nascent and mature RNA expression noise. We re-ran the two linear regressions shown in figures 3 and 4B, but incorporating log_2_(miRNA binding site count) as the second predictive variable. The result is schematically shown in figure 6B. The regression analysis revealed a clear trend that mature RNAs with higher binding site counts exhibited lower expression noise compared to nascent RNAs (Fig. 6B). ANOVA comparison of the regression models determined that adding the second predictive variable led to significant improvement (F_1_ = 133, p-value < 2.2E-16).

Thus, our combination of mathematical modeling and snapTotal-seq data analysis confirmed that miRNA-targeted mRNAs should have reduced inter-cell expression noise. To the best of our knowledge, this is the first report of using total-RNA scRNA-seq data to investigate miRNA-targeted mRNA degradation and expression noise reduction simultaneously. Next, we delved into the functional implications of our previous observation that major components of miRNA-mediated regulatory pathway themselves are top miRNA targets^21,22^.

### The miRISC negative self-feedback loop

We previously uncovered the AGO and TNRC6 negative self-feedback loops while analyzing miRNA binding site distribution^21,22^. Briefly, when human mRNAs were ranked in descending order by their unique miRNA binding site counts, AGO1/2/3 and TNRC6A/B/C are all highly ranked. AGO1 is ranked at the 5^th^, AGO2 the 6th, AGO3 the 11^th^ (tied) and TNRC6B the 27^th^; TNRC6C and TNRC6A are respectively ranked, though not as high but both within the top 4%, at the 446^th^ and the 800^th^.

Additionally, the RNA binding protein QKI, whose mRNA has 62 miRNA binding sites and is ranked at the 15^th^ among all human mRNAs, was recently reported as an auxiliary partner for miRNA-AGO1/2/3 regulatory activity^38^. It binds to both mature miRNAs and AGO proteins, contributing to miRNA stabilization and target mRNA decays^38,45,46^. Thus, QKI is an auxiliary participant in the miRISC negative self-feedback loop. However, it is not the only QKI feedback loop. QKI also forms a negative feedback loop with the E2F1 transcription factor to control its transcription; E2F1 activates QKI transcription, and QKI in turn regulate the RNAs of E2F1 and its partners to suppresses E2F1 activity^47^.

This dataset gave us an opportunity to assess the function of the miRISC self-feedback loop, as AGO1/2/3, TNRC6A/B/C and QKI mRNAs are all detected in every cells. As discussed in the Materials and Methods section, we mathematically modeled the AGO/TNRC6 negative self-feedback loop as adding a stabilizing element (*σ*R) to K_deg_. When higher-than-intended expression levels of a feedback-controlled mRNA leads to higher protein expression levels, the feedback path would channel higher miRNA-mediated degradation activity back onto the same mRNA, reducing expression levels back to normal level (R*). And the opposite would happen when the controlled mRNA is expressed at lower-than-expected levels. Impacts of the feedback element (*a*R) on S and CV are predicted below:

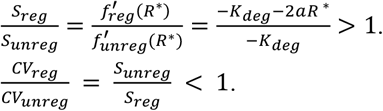

Thus, the self-feedback loop should further reduce the inter-cell expression noise (CV) of controlled mRNAs.

To test this prediction, we re-plotted figure 2C and highlighted relevant mRNAs for comparative analyses (Fig. 7). The 16 mRNAs with 60 or more miRNA binding sites that were detected in every cell were highlighted in red (AGO1/2/3), purple (QKI) and blue (others) colors. The TNRC6A/B/C mRNAs are highlighted in green (Fig. 7). The dotted curves show the variation of expected residuals/errors above and below the weighted linear regression line (Fig. 7), facilitating visual comparison of the CVs of mRNAs with different expression levels. As expected, AGO1/2/3 mRNAs all have low CVs in comparison to other mRNAs with 60 or more conserved miRNA binding sites and similar mean expression levels. Consistent with its feedback control at both transcription and mRNA degradation (K_deg_) steps, QKI has even lower CV. TNRC6A/B/C mRNAs, though having less than 60 binding sites, also have relatively low noise levels (Fig. 7). Thus, miRNA-targeted mRNAs regulated by negative self-feedback loops have further reduced inter-cell expression noise.

**Figure 7.**
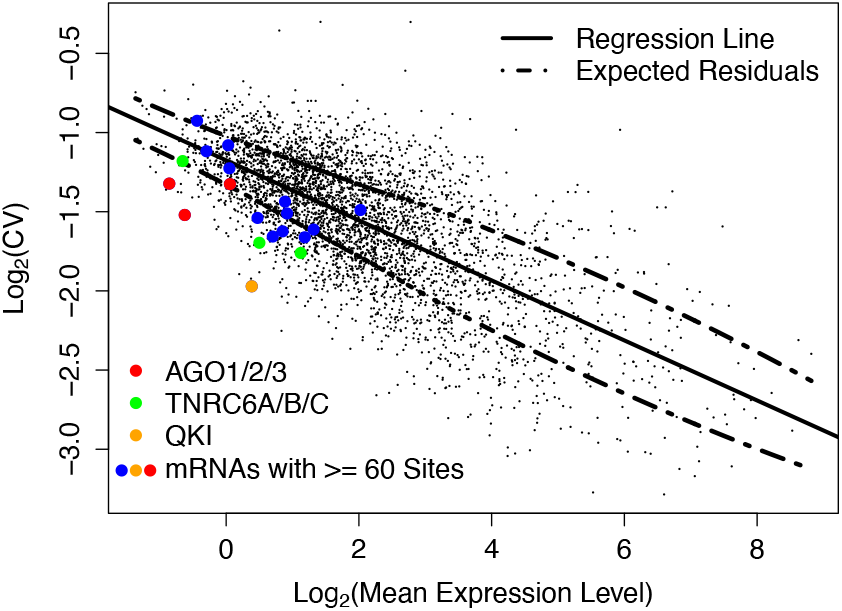
Reduced expression levels and noises of miRISC mRNAs. Re-plot of figure 2C, high-lighting mRNAs with >= 60 miRNA binding sites as red (AGO1/2/3), orange (QKI, an auxiliary miRISC component) and blue (others) data points. TNRC6A/B/C are highlighted as green data points. Dashed curves show the variation of expected residuals above and below the regression line.

Statistical analysis confirmed this pattern quantitatively. The weighted residuals of the linear regression were used as expression-level-adjusted mRNA CVs. A t-test showed the four feedback-controlled mRNAs with more than 60 miRNA binding sites (AGO1/2/3 and QKI) have lower adjusted CV values than the other 12 mRNAs with 60 or more binding sites (T_14_ = −3.34 and P-value = 0.0024). When QKI was not included in the comparison, AGO1/2/3 mRNAs alone showed significant lower values than the other 12 mRNAs (T_13_ = −2.41 and P-value = 0.016). Despite having less than 60 binding sites, TNRC6A/B/C mRNAs also showed significant lower values than the 12 mRNAs (T_13_ = −1.96 and P-value = 0.036).

### Confirmation that miRNAs exert their regulatory activities at the mature mRNA step

Having the expression levels at nascent and mature mRNA steps simultaneously gave us an opportunity to determine the step, at which gene expression noise reduction occurs for individual genes. As discussed above, we already observed reduced expression levels of miRNA-targeted mRNAs (Fig. 1C), but not for their counterparts at the nascent RNA step (Fig. 1B). We investigated whether this pattern was also applicable for gene expression noise reduction, by re-plotting figure 2D and highlighting the counterparts for miRNA-targeted mRNAs (with >= 60 binding sites) as red (AGO1/2/3), orange (QKI) and blue (others) data points (Fig. 8). As expected, the nascent-RNA counterparts exhibit normal gene expression noise levels (Fig. 8). And the pattern is generic to the whole transcriptome (Fig. 5B). The trends of decreasing expression levels and inter-cell noise as miRNA binding site count increases observed at the mature RNA step (Fig. 5A) were absent at the nascent RNA step (Fig. 5B). The results demonstrated the power of total-RNA scRNA-seq in studying miRNA and other post-transcriptional regulatory mechanisms.

**Figure 8.**
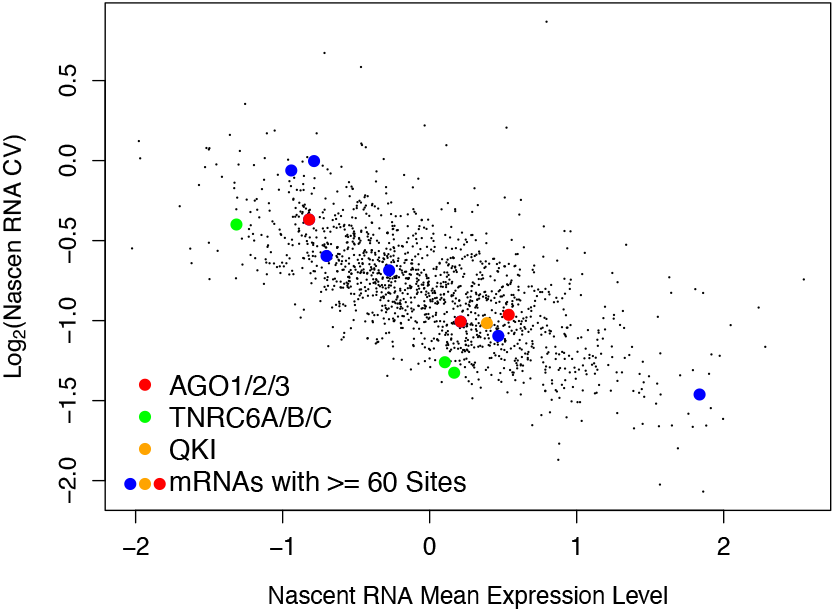
Scatter plot of nascent RNA expression noise versus expression level. Counterparts of mRNAs with >= 60 miRNA binding sites are highlighted as red (AGO1/2/3), orange (QKI) and blue (others) data points. And TNRC6A/B/C are highlighted as green data points.

### Differential control of AGO1/2/3, TNRC6A/B/C and QKI RNA inter-cell expression noise

Next, we focused on the AGO1/2/3, TNRC6A/B/C and QKI RNAs. As other mRNAs with 60 or more conserved miRNA binding sites, AGO1/2/3 and QKI mRNAs are controlled at the mature mRNA step. They did not show highly reduced expression levels and noises at the nascent RNA step (Fig. 8) but did so at the mature mRNA step (Fig. 7). However, TNRC6A/B/C already exhibited low inter-cell expression noise levels at the nascent RNA step (Fig. 8). The self-feedback loop reinforces the low noise levels at the mature mRNA step (Fig. 7).

## Discussion

The importance of the miRNA-mediated gene expression regulation subsystem is underscored by, among many others, its evolutionary conservation^1-4^, pathogenic mutations in diseases^48-50^ and the recent award of the 2024 Nobel Prize in Physiology or Medicine to its discovery^51^. While entering this research area, we have expressed curiosity on why this subsystem is maintained so ubiquitously, despite that it seems metabolically and energetically wasteful^21,50,52^. Via the cell-computer analogy that we previously examined^53,54^, we speculated its role in alleviation of operational latency caused by the time delay from transcription to translation.

One of the key functional advantages attributed to the miRNA regulatory mechanism is the enhancement of gene expression level stability. In this study, we took advantage of a total RNA scRNA-seq dataset, to the best of our knowledge for the first time, to simultaneously examine miRNA-mediated mRNA degradation and expression noise reduction. Although our analysis is limited to the mRNA step, it can be argued that the reduction in inter-cell mRNA expression noises is propagated to downstream steps, leading to reduced expression noise at the translation and protein steps.

This gain of operational advantage, in exchange for maintaining the miRNA-mediated gene expression regulatory subsystem, aligns with the evolutionary co-emergence of this subsystem with multicellularity, that is, its ubiquitous occurrence in such species. Organismal-level functionalities and processes of multicellular species depend on cellular homogeneity and robust dynamic state transitions, defectives of which are often pathogenic. At the cellular level, stable steady-state gene expression level is central to cellular homogeneity. For instance, as discussed in the Introduction section, a major hallmark of cancer is the loss of this homogeneity to enable cancer cell clonal growth and metastasis. Not surprisingly, miRNAs have been shown to be globally depleted in cancers^55^. Furthermore, the DICER1 syndrome, caused by a heterozygous mutation of the DICER1 gene, is associated with elevated cancer risks^56^. Thus, to optimize multicellular operations, such organisms maintain the miRNA-mediated regulatory subsystem at the expense of the seemingly wasteful metabolic and energetic expenditure to produce miRNA-targeted mRNAs.

Our results also supported the noise reduction function of the miRISC negative self-feedback regulation loops. The AGO and the TNRC6 proteins form the miRNA targeting apparatus and the core of the miRISC complex, and their mRNAs are controlled by self-feedback loops^21,22^. The AGO proteins process the double-stranded precursors, which is passed onto them from DICER1, into mature miRNAs and orient them for target binding^57^. Upon targeting binding, the miRNA-AGO complex recruits the TNRC6 proteins, which subsequently recruit downstream effecter proteins^58^. Thus, AGO and TNRC6 channel the miRNA-mediated regulatory actions on the target mRNAs. Since AGO and TNRC6 mRNAs themselves are top miRNA targets, they channel significant regulatory actions back onto their own mRNAs to form close negative self-feedback loops. We observed that the inter-cell expression noise of AGO1/2/3 and TNRC6A/B/C mRNAs was further reduced, which is consistent with the feedback loop acting as a common noise reduction mechanism.

To the best of our knowledge, the miRISC negative self-feedback loop is the first that adds the feedback regulatory element onto mRNA degradation activity (K_deg_). Most reported gene expression feedback loops add the regulatory elements onto transcription activity, and so are most of the loops in engineered regulatory circuits in synthetic biology. One exception is the feedback control of the nuclear export of its own nascent RNA by the HIV Rev protein, which reduces mRNA expression noise and is a critical HIV pathogenesis factor^59^. Additionally, the ribonucleoprotein particles (RNP) often feedback control the mRNAs for their own proteins^60^. However, none of feedback loops have been shown to directly add the regulatory element to mRNA K_deg_. Nevertheless, the feedback control appears to be a ubiquitous regulatory mechanism throughout the entire gene expression process.

The differential control of AGO and TNRC6 mRNA inter-cell expression noise – AGOs at the mature mRNA step and TNRC6s at the nascent RNA step – is intriguing. Conceivably, it is related to the functional difference between the AGOs and the TNRC6s. Even though both are major miRISC components, they do have differences in other functional aspects. For instance, in addition to being part of the miRISC, TNRC6 proteins are considered the p-body scaffold proteins^61^, while AGO proteins are not. Multiple RNA-silencing pathways converge on TNRC6^62^. Additionally, AGO1/2/3 proteins are well-known to be associated with ribosomes (both monosomes and polysomes)^63^; they were used to be also known/named as eukaryotic translation initiation factor 2Cs, EIF2C1/2/3. To the best of our knowledge, TNRC6 proteins have not been shown to do so.

However, another total RNA scRNA-seq dataset that traces the cells as they go through a dynamic physiological process is necessary to investigate another functional aspect of the self-feedback loop – ensuring smooth dynamic state transitions. In this context, a transition means a shift from one steady-state expression level to another. For example, when the transcription rate of a gene increases while the cognate mRNA K_deg_ remains the same, the mRNA steady-state expression level should increase accordingly. According to the control theory, a feedback loop should help to ensure a smooth transition process^27^. A dataset covering a cell population at a single condition or time-point is not sufficient for analyzing such dynamic processes.

In summary, this study took advantage of a total-RNA scRNA-seq dataset, whose NGS sequencing chemistry covers full-length RNAs. This enabled higher detection sensitivity, more than 10,000 genes in each cell, at both the nascent RNA and the mature mRNA steps of the gene expression process. We were able to investigate simultaneously the miRNA-mediated target mRNA degradation and inter-cell expression noise reduction, as well as the function of the miRISC negative self-feedback loop. This study demonstrated the power of the total RNA scRNA-seq technologies for such studies, paving the ground for comparative study of wild-type and isogenic mutant cells as well as cells at different stages of dynamic physiological processes.

## Acknowledgement

We acknowledge Mr. Emmanuel Annan for participation in scientific discussion.

